# TSG Targeting KDM5A affects osteogenic differentiation of bone mesenchymal stem cells induced by bone morphogenetic protein 2

**DOI:** 10.1101/2020.07.25.221424

**Authors:** Min Wei, Yi Jiang, Yuanqing Huang

## Abstract

To investigate the effect of 2, 3, 5, 4’-tetrahydroxystilbene-2-O-β-D-glucoside (TSG) on osteogenic differentiation of bone marrow mesenchymal stem cells (BMSC) and its molecular mechanism. The effects of TSG on alkaline phosphatase positive cloning and mineralized nodule formation were also detected. Total mRNA and protein were extracted and effects of TSG on the expression levels of osteopontin (OPN), osteocalcin (OCN), Runt-related transcription factor 2 (Runx2), Osterix and Col1a1 were detected by real-time fluorescence quantitative PCR. Western Blotting was used to detect the inhibitory effect of TSG on KDM5A. BMSCs were transfected with Small interfering RNA (siRNA) targeting KDM5A (si-KMD5A) and pcDNA3.1 KMD5A. TSG significantly increased the activity of ALP and the number of alkaline phosphatase clones and calcified nodule formation. The OPN, OCN, Runx2 and Osterix expression levels were significantly increased among the osteoblasts after TSG treatment. The mechanism study showed that the effect of TSG is realized by inhibiting KDM5A. KDM5A signaling may be involved in the regulation of osteogenic differentiation of rBMSC. TSG can promote osteogenic differentiation and maturation of rBMSC at 0.1-50 μmol / L. The mechanism of action was realized by inhibiting the expression of KDM5A.

## Introduction

Osteoporosis (OP) is a metabolic bone disease characterized by reduced bone mass, damaged bone mass and reduced bone strength, resulting in increased bone fragility and prone to fracture. Osteoporosis is characterized by complexity, comprehensiveness, youth and masculinity. Therefore, the research on the etiology and pathogenesis of osteoporosis must have a new understanding and improvement. Bone marrow mesenchymal stem cells (BMSCs) are derived from bone marrow and have the function of self-renewal and multidifferentiation(Guan et al., 2012). The osteogenic differentiation ability of BMSCs decreases with age(Baker, Boyette, & Tuan, 2015; ZHANG, LU, & SUN, 2014). In osteoporosis patients, the proliferation and self-renewal capacity of MSCs are reduced, the number of apoptosis is increased, osteogenic differentiation is weakened, lipogenic differentiation is enhanced, and the regulation of multidirectional differentiation is disordered(Carbonare et al., 2009; Muruganandan, Roman, Sinal, & sciences, 2009; Zhou et al., 2008).

TSG is an active component of Polygonum multiflorum and has a wide range of biological activities(Lin et al., 2017; Xia, Wei-fen, & Yan-jun, 2012; Xiang et al., 2014). Studies have shown that TSG has anti-aging, anti-atherosclerosis, anti-hyperlipidemia, anti-tumor, anti-inflammation, free radical scavenging, liver protection and other biological activities(Huang et al., 2013; Long, Zhang, Gao, Qiao, & Tian, 2011; Shi-Yin et al., 2011; Song et al., 2015; X. P. Yang, Liu, Qin, & Yu, 2014). TSG has a wide application prospect.

Histone methylation modification is one of the most important aspects of epigenetic research. Histone methylation is dynamically regulated by histone methylase and histone demethylase. Histone lysine demethylase 5A (KDM5A) is an important member of the histone demethylase family. KDM5A can specifically remove dimethyl and trimethyl (H3K4me2/3) from the fourth lysine of histone H3, thereby mediating gene silencing and regulating cell function(Boschiero et al., 2013; Hou, Wu, Dombkowski, Zhang, & Yang, 2012). KDM5A can directly or indirectly maintain the tumor cell dryness, inhibit cell metabolism and differentiation, and promote the proliferation, metastasis and drug resistance of tumor cells. Therefore, KDM5A is closely related to the occurrence and development of a variety of diseases and is expected to be a new potential target for the treatment of osteoporosis.

In this study, the effects of TSG on osteogenic differentiation and maturation as well as the regulation of KDM5A signaling pathway were studied in rat rBMSC. To provide potential targets and candidate compounds for the treatment of osteoporosis.

## Results

### Effects of TSG on the toxicity and osteogenic differentiation of bone marrow mesenchymal stem cells (rBMSCs)

Fig. 1A shows the molecular structure formula of TSG. To investigate effects of TSG on BMSCS, we first analyzed the effects of different concentrations of TSG on cell proliferation. Experimental results showed that TSG did not affect cell proliferation (Fig. 1B). Further, we evaluated the effect of TSG on alkaline phosphatase in rBMSC cells. Experimental results showed that alkaline phosphatase activity positive clone (CFU-FALP) could be formed in both the control group and administration group. After TSG treatment (1, 10 and 50μmol/L), the area, quantity and relative gray level of CFU-FALP were higher than those of the control group, and the difference between the two groups was statistically significant (Fig. 1C). Further experimental results showed that different concentrations of TSG increased ALP activity (Fig. 1D) and presented a concentration-dependent effect. The influence of TSG on the formation of calcified nodules the results showed that both the control group and the TSG group formed calcified nodules after osteogenic induction of rBMSC. After TSG treatment (1, 10 and 50μmol/L), the number of calcified nodules was more than that of the control group. According to IPP 6.0 software analyses, the area, number and relative gray scale of calcified nodules were higher than those of the control group, and the difference between the two groups was significant (Fig. 1E,1F). Further, we examined the effect of TSG on osteogenic differentiation related proteins. The results showed that TSG treatment with different concentrations could up-regulate the expression of OPN, OCN, Runx2 and Osterix (Fig. 2A-2D).

**Figure 1.**
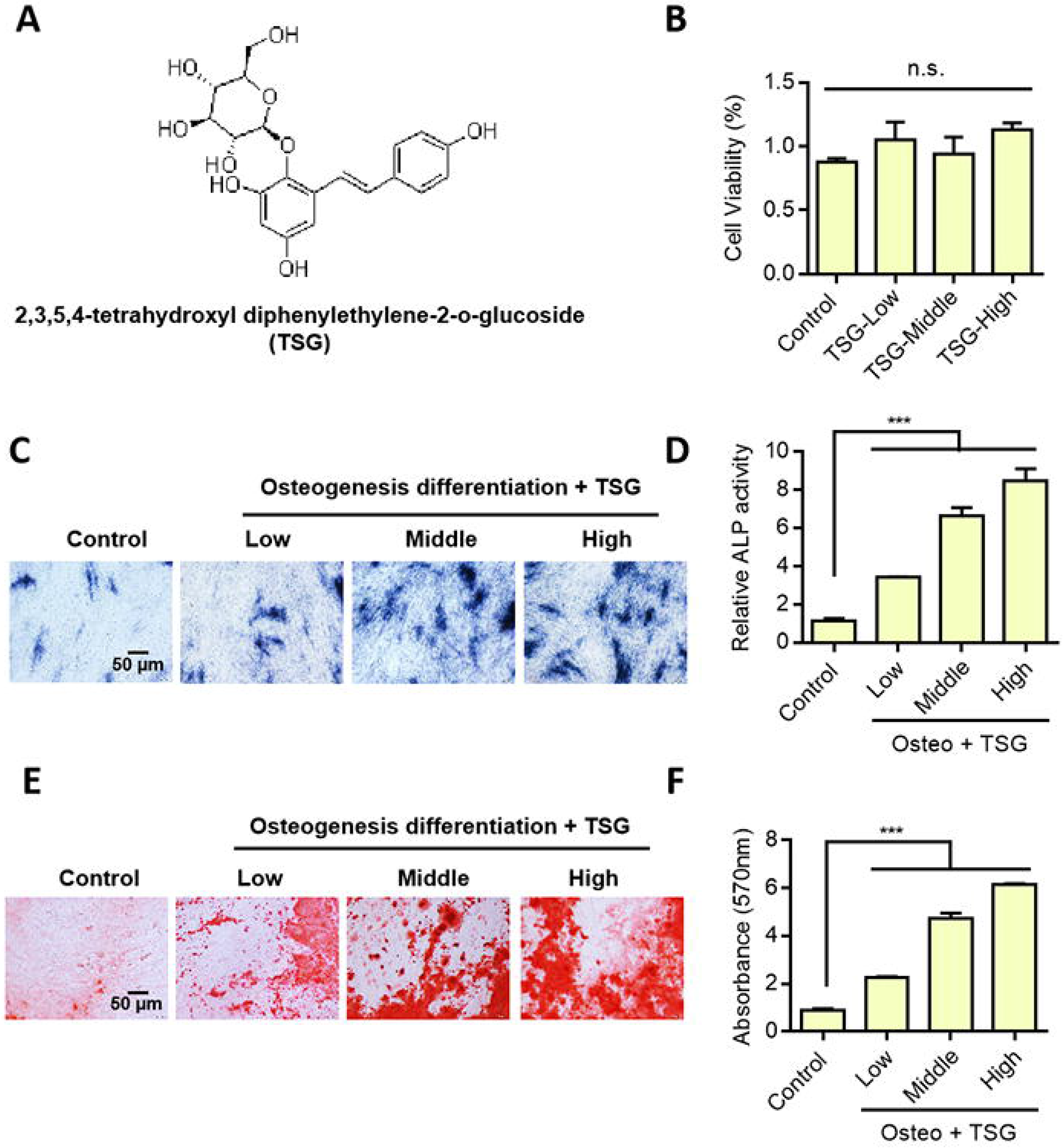
Effects of TSG on the toxicity and osteogenic differentiation of bone marrow mesenchymal stem cells (rBMSCs). A. Display of TSG molecular structure. B. RBMSCs were treated with TSG of different concentrations for 1 ~ 2 days, and cell count CCK-8 was used to detect cytotoxic effects. C. TSG treatment of rBMSCs for 1 week, and Alkaline phosphatase staining of rBMSCs with D. TSG treatment for 1 week, the activity of Alkaline phosphatase was detected. E. Mineralization of rBMSCs 2 weeks after TSG staining detection. F. Statistical results of Mineralization of rBMSCs 2 weeks after Alizarin Red S staining detection TSG staining detection. Data were mean ± S.D, \*\*\**P*<0.001.

**Figure 2.**
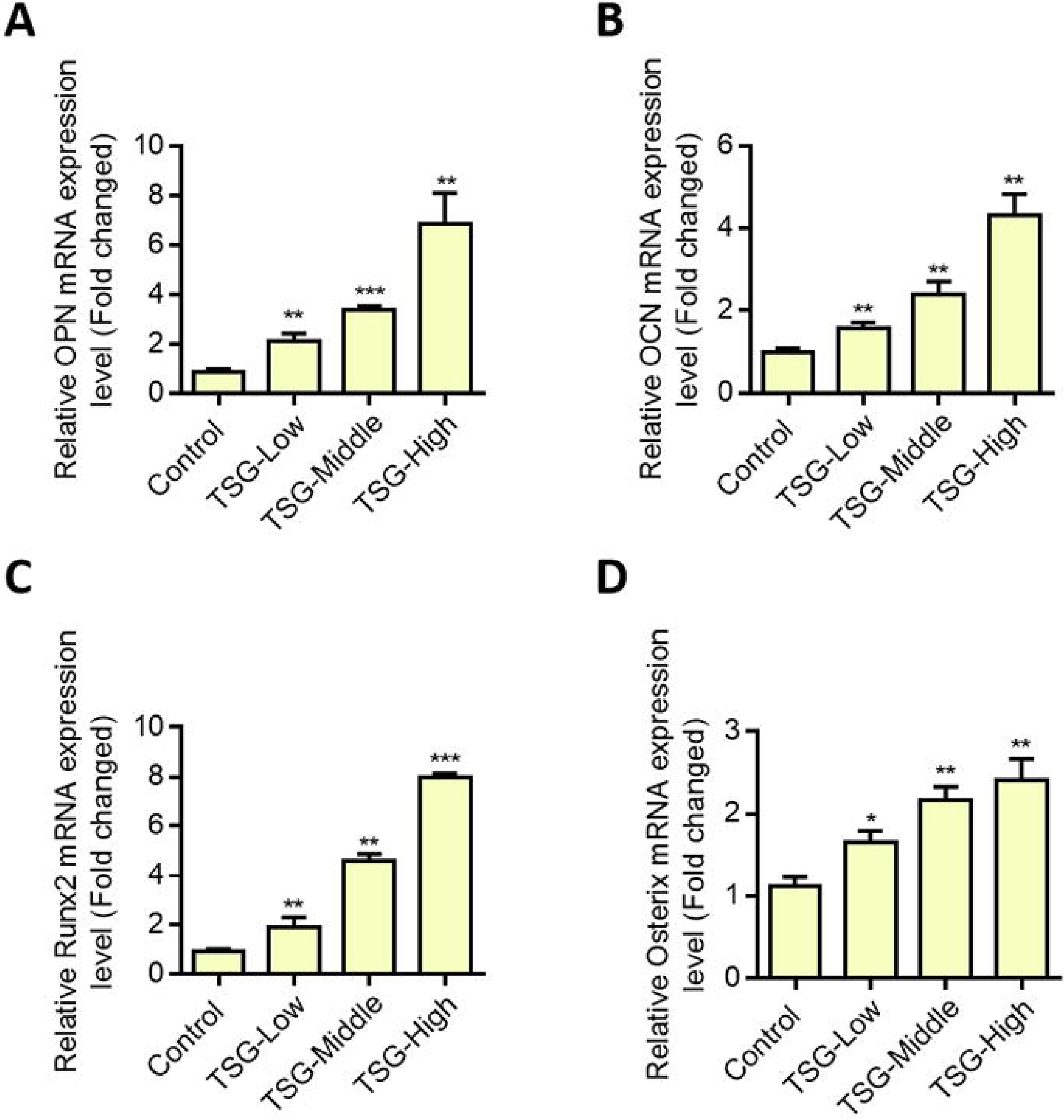
Effects of TSG on the expression of osteopontin (OPN), osteocalcin (OCN), Runt-related transcription factor 2 (Runx2) and Osterix. A. Detection of OPN expression by qPCR. B. OCN expression was detected by qPCR. C. QPCR was used to detect Runx2 expression. D. Detection of Osterix expression by qPCR. Data were mean ± S.D, \**P*<0.05, \*\**P*<0.01, \*\*\**P*<0.001.

### Knockdown KDM5A promoted osteogenic differentiation of bone marrow mesenchymal stem cells

Through experiments, we found that the expression of KDM5A was decreased during the osteogenic differentiation induced by BMP-2 (Fig. 3A). Western blot results also showed that bMP-2 induction could inhibit the expression of KDM5A (Fig. 3B). To further investigate the direct role of KDM5A in osteogenesis of MSCs, we knocked down KDM5A in normal MSCs (Fig. 4A). Alkaline phosphatase staining results showed that knockdown KDM5A could promote ALP activity under the osteogenic differentiation induced by BMP-2 (Fig. 4B). Alizarin red staining showed the knockdown of KDM5A, which significantly promoted the bMP2-induced mineralization deposition (Fig. 4C). In addition, RT-PCR showed that knockdown KDM5A could up-regulate the expression levels of OPN, OCN, Runx2, Osterix and Col1a1 (Fig. 4D-4H). These results indicate that KDM5A negatively regulates osteoblast differentiation. Elevated KDM5A may be the cause of osteoporosis.

**Figure 3.**
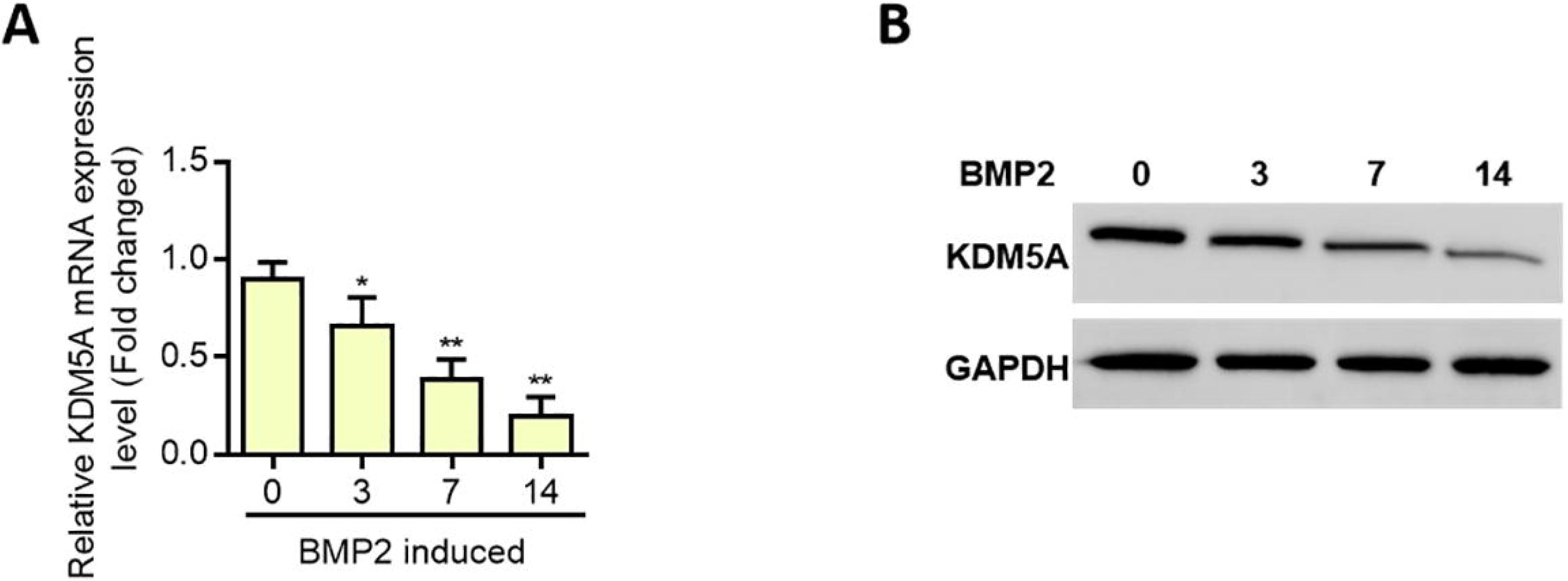
KDM5A expression was down-regulated under osteogenic differentiation induced by bone morphogenetic protein 2 (BMP-2). A. The expression level of KDM5A was detected by qPCR. B. Western blot detection of KDM5A expression. Data were mean ± S.D, *P<0.05, **P<0.01.

**Figure 4.**
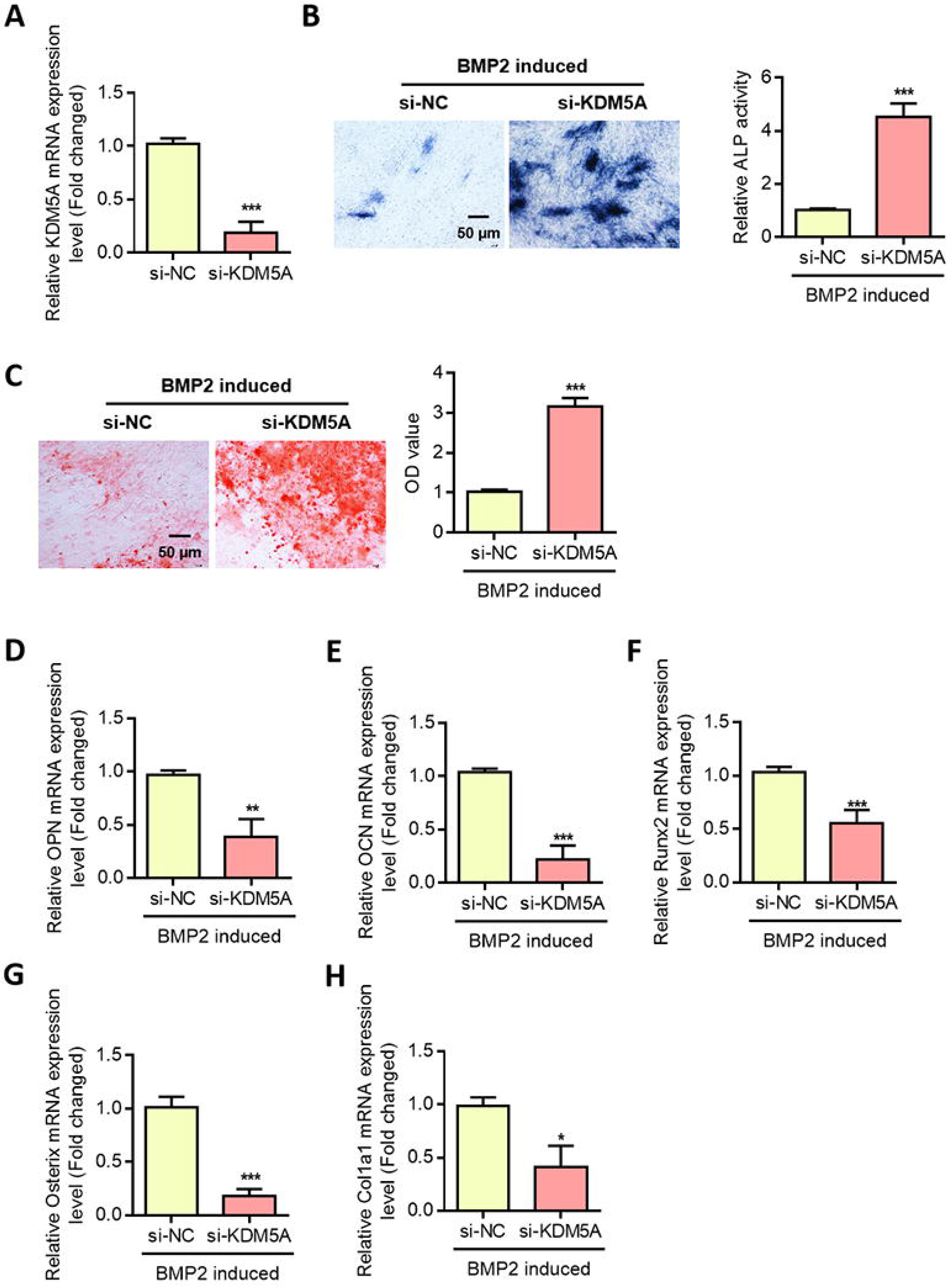
Knockout KDM5A promoted osteogenic differentiation of bone marrow mesenchymal stem cells. A. QRT-PCR analysis of KDM5A expression in transfected MSCs. B. Alkaline phosphatase staining and activity of MSCs with si-KDM5A were detected by ALP staining 7 days after osteogenesis induction. C. Use Alizarin Red S staining to detect mineralized nodules of bone marrow mesenchymal stem cells after si-KDM5A. D. QPCR was used to detect OPN expression in MSCs of si-KDM5A after osteogenesis induction. E. qPCR detection of OCN expression in MSCs of si-KDM5A after osteogenesis induction. F. QPCR was used to detect Runx2 expression in MSCs of si-KDM5A after osteogenesis induction. G. QPCR was used to detect Osterix expression in MSCs of si-KDM5A after osteogenesis induction. H. QPCR was used to detect Col1a1 expression in MSCs of si-KDM5A after osteogenesis induction. Data were mean ± S.D, \**P*<0.05, \*\**P*<0.01, \*\*\**P*<0.001.

### Influence of TSG on KDM5A expression

Further, we evaluated the effect of TSG on KDM5A expression. First, we treated primary rBMSC with different concentrations of TSG. By real time-qPCR detection KDM5A expression changes, with a relative quantitative method 2^−ΔΔCT^ analysis of test results. The experimental results showed that compared with the control group, the gene expression of KDM5A was significantly inhibited in the TSG group with different concentrations (Fig. 5A). Western blot results were consistent with qRT-PCR results. Experimental results showed that TSG treatment could inhibit the protein expression of KDM5A (Fig. 5B).

**Figure 5.**
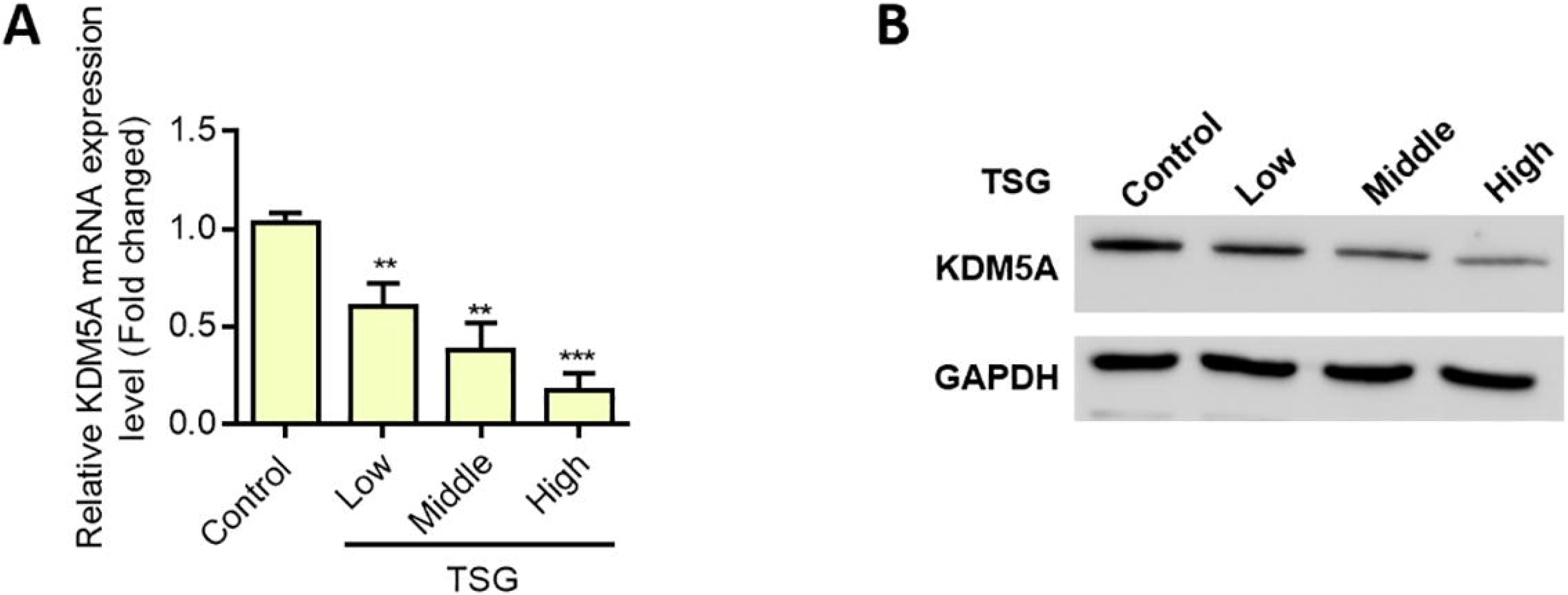
TSG treatment inhibits the expression of KDM5A. A. The expression level of KDM5A was detected by qPCR. B. Western blot detection of KDM5A expression. Data were mean ± S.D, \*\**P*<0.01, \*\*\**P*<0.001.

### TSG promotes osteogenic differentiation of bone mesenchymal stem cells by targeting the expression of KDM5A

In order to further investigate the direct effect of TSG on osteogenesis of MSCs by inhibiting KDM5A, we overexpressed KDM5A in normal MSCs (Fig. 6A). Alkaline phosphatase staining results showed that under the osteogenic differentiation induced by BMP-2, overexpression of KDM5A could inhibit the activity of ALP, while TSG treatment could up-regulate the activity of ALP (Fig. 6B). Alizarin red staining showed that overexpression of KDM5A significantly reduced BMP2-induced mineralization deposition, while TSG treatment was able to up-regulate mineralization (Fig. 6C). In addition, RT-PCR showed that KDM5A overexpression could inhibit the expression of OPN, OCN, Runx2, Osterix and Col1a1, while TSG treatment could up-regulate the expression of these genes (Fig. 6D-6H). These results indicated that KDM5A inhibited osteoblast differentiation, while TSG could play a protective role by inhibiting the expression of KDM5A.

**Figure 6.**
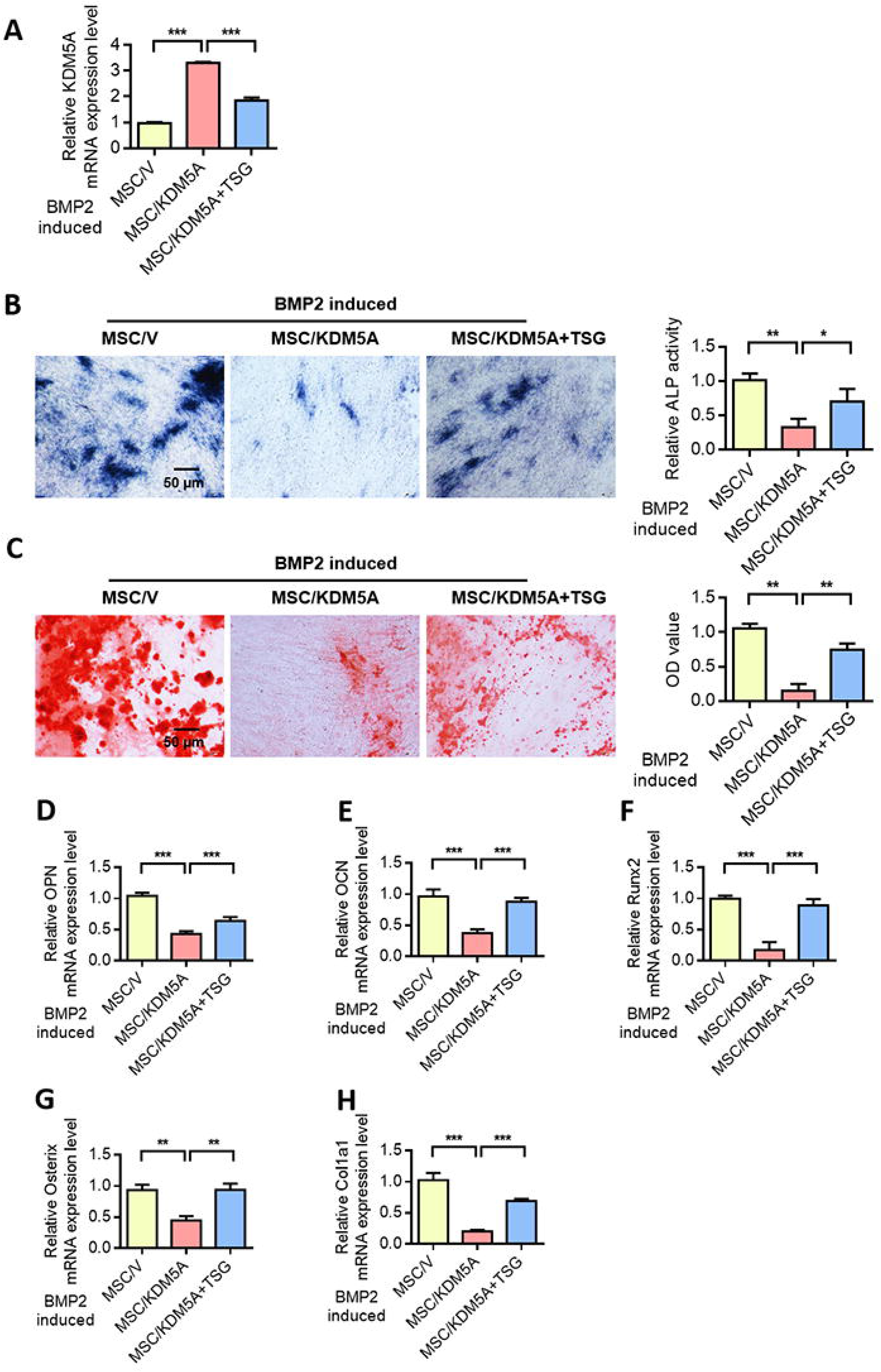
TSG promotes osteogenic differentiation of bone mesenchymal stem cells by targeting the expression of KDM5A. A. QRT-PCR analysis of KDM5A expression in MSCs after different treatments. B. Alkaline phosphatase activity was detected by ALP staining after osteogenic induction of MSCs in each group. C. Use Alizarin Red S staining detection for mineralized nodules formed by bone marrow mesenchymal stem cells after staining detection in each group. D. QPCR was used to detect the OPN expression of MSCs in each group after osteogenesis induction. E. QPCR detection of OCN expression in MSCs treated in each group after osteogenesis induction. F. QPCR was used to detect Runx2 expression of MSCs in each group after osteogenic induction. G. QPCR was used to detect Osterix expression in MSCs treated in each group after osteogenic induction. H. QPCR detected the expression level of Col1a1 of MSCs in each group after osteogenic induction. Data were mean ± S.D, \**P*<0.05, \*\**P*<0.01, \*\*\**P*<0.001.

## Discussion

BMSCs are derived from bone marrow and have the functions of self-renewal and multidifferentiation. It can differentiate into osteoblasts, adipocytes, myoblast, chondroblast and nerve cells under certain conditions(Guan et al., 2012). BMSCs can promote osteogenic differentiation and increase bone formation and strength. In recent years, the application of BMSCs in cell therapy of bone-related diseases has attracted great attention. BMSCs also show great application potential in bone tissue engineering(Bianco et al., 2013; Guan et al., 2012; Olivares-Navarrete et al., 2015). The results showed that TSG could promote osteogenic differentiation and maturation of BMSCs, significantly improve ALP activity, ALP positive clone formation and calcified nodule formation. TSG can increase the expression of Runx2 and other genes, which are closely related to the osteogenic differentiation of cells. In recent years, studies have shown that plant drugs have great potential in the prevention and treatment of osteoporosis. On the basis of traditional Chinese medicine formulations, several Chinese patent medicines against osteoporosis have been developed and improved. Wang et al. (Wang et al., 2013) summarized 12 randomized controlled trials of proprietary Anti-osteoporosis drugs and evaluated their effectiveness, including 1816 patients with osteoporosis. The results showed that all the herbal preparations significantly increased the lumbar spine bone density. A variety of flavonoid compounds have been shown to contribute significantly to bone formation at the cellular or animal levels. For example, icariin, genistein, resveratrol, osthole, etc(Durbin et al., 2012; Wang et al., 2013; L. Yang, Yu, Qu, Li, & biophysics, 2014; Zhai, Guo, et al., 2014; Zhai, Pan, et al., 2014). Compared with traditional osteoporosis medicine, plant medicine has wide sources and low side effects(Su-Jin et al., 2018; Zhao et al., 2013). Therefore, it has great application development value. TSG is the main component of Polygonum multiflorum and has good water solubility(Song et al., 2015; X. P. Yang et al., 2014). At the same time, it has the function of the antioxidant and scavenging free radicals. TSG also has a variety of biological regulatory effects, such as anti-osteoporosis, vasodilation, anti-complement activity, etc. In this study, it was found that TSG promoted osteoblast differentiation not through estrogen signaling pathway, but by inhibiting the expression of KDM5A. We found that TSG can enhance the expression of OPN, OCN, Runx2 and Osterix genes. Western blot further proved that TSG treatment could reduce the protein expression level of KDM5A. The results show that TSG may promote the differentiation and maturation of BMSCs by regulating KDM5A signals. KDM5A is a histone demethylase that can specifically remove dimethyl and trimethyl (H3K4me2/me3) from the fourth lysine of histone H3, so it is also known as H3K4me2/3 demethylase(Dezhi et al., 2016; Nishibuchi et al., 2014; Torres et al., 2015). KDM5A plays an important role in cell development and differentiation(Dahl et al., 2016). At present, the regulation mechanism of KDM5A on cell differentiation is not very clear. Several studies have shown that KDM5A regulates cell differentiation mainly through indirect pathways. In mouse embryonic stem cells, KDM5A can synergistic effect with Krüppel-like factor 4 (KLF4) to block the programming of pluripotent stem cell differentiation(Dabiri et al., 2019). Furthermore, KDM5A maintained the desmethylation state of core pluripotent transcription factor POU5F1, thereby inhibiting embryonic stem cell differentiation and maintaining cell dricity 15. In addition, KDM5A can also form a co-inhibitory complex with polycomb group (PcG) to jointly maintain cell driness and inhibit cell differentiation(Pasini et al., 2008).

This study also has deficiencies. The differentiation and maturation of BMSCs was a complex process, and whether other signaling molecules or other signaling pathways are involved in the TSG regulation of osteogenic differentiation of rBMSC remains to be further studied.

In conclusion, KDM5A is involved in the regulation of osteogenic differentiation of MSCs. Inhibition of KDM5A promoted osteogenic differentiation of MSCs. TSG was added in MSCs culture to promote differentiation and mineralization to osteoblasts without affecting cell proliferation. Experimental results confirmed that TSG can promote osteogenic differentiation. Further research on the specific mechanism of TSG and KDM5A in regulating MSCs cell behavior will provide the experimental basis for gene targeted therapy and drug cell therapy for OP.

## Methods

### Isolation and culture of rat bone marrow mesenchymal stem cells

5 rats (12 weeks old) were obtained from Vital River Laboratory Animal Technology Co., Ltd. (Beijing, China) and were anesthetized by intraperitoneal injection of 10 mg / kg phenobarbital sodium. And the rats were killed by CO_2_ suffocation. Immerse in 75% alcohol for 10 min. Femur and tibia were rapidly separated under aseptic conditions and epiphyses were removed at both ends. DMEM/F12 culture medium was extracted with a 5 mL syringe to flush the bone marrow cavity until the liquid was clarified. The rinse solution was collected, blown with a micropipette and left to stand for 5 min. The supernatant was centrifuged at 1000 r / min for 5 min and the supernatant was discarded. The cells were resuspended with DMEM/F12 containing 10% FBS. After repeated blowing and counting, 10^4^ cells / cm^2^ were inoculated into a 6-well plate. Culture in an incubator at 37 ◻ and 5% CO_2_. First liquid changed after 3 days. Cell growth was observed daily. After 80% of the adherent cells were fused, trypsin was used for digestion and subculture (this was the first generation P1). When cells were passed to P3, the inoculation densities of 96-well plate, 24-well plate and 6-well plate were 2×10^3^, 2×10^5^ and 4×10^5^ cells / well, respectively. All experiments were performed in Hunan University of Medicine and got approval of animal ethics committee of Hunan University of Medicine.

### Induction of bone differentiation

When the P3 generation MSCs in each group reached 80% ~ 90% fusion, the digestive cells were digested. MSCs were inoculated into a six-well plate. After 24 h, the old culture medium was removed and 2 mL of osteogenic induction liquid (bone morphogenetic protein 2) was added to each well. Liquid was exchanged every 3 days for 21 consecutive days. The treatment group was given TSG at concentrations of 1, 10 and 50 μmol / L.

### Cell transfection

The small interfering RNA (siRNA) targeting KDM5A (si-KMD5A), siRNA negative control (si-NC) were all obtained from Shanghai GenePharma Co., Ltd. (Shanghai, China). The cDNA encoding KMD5A was amplifified by PCR, and then ligated into pcDNA3.1 (+) vector (Invitrogen). Cell transfection was performed using Lipofectamine 2000 (Invitrogen). After 48 h, the transfection effificiency was validated by RT-qPCR analysis.

### Determination of alkaline phosphatase activity

The cells were inoculated in 12-well plates and cultured for 24 h for osteogenesis induction. The treatment group was given 10 μmol / L TSG, and the control group was given the blank induction medium. The culture medium of both the administration group and the control group contained 1‰ DMSO. After 7 days, ALP histochemical staining was performed by azo method. Wash with PBS twice, fix with 10% formaldehyde solution for 1min, add matrix solution (Michaelis barbiturate-HCL buffer containing 0.1% α-naphthol sodium phosphate and solid blue RR salt, pH 8.9) and react with 30 min. When brown spots appeared, the matrix solution was discarded, washed with PBS and fixed, and the results were photographed for preservation. The area, quantity and gray level of CFU-FALP was scanned by image-Proplus6.0 software for statistical analysis.

### Alizarin Red S staining

The cells were inoculated in a 24-well plate. Osteogenic differentiation was induced after 24 h culture. The administration group was given 10μmol/L TSG. The blank induction medium was added to the control group, and the culture medium in both the administration group and the control group contained 1‰ DMSO. Discard the culture medium on day 12. Wash with PBS twice, and fix with 10% formaldehyde for 5min. Alizarin red dye was added and incubated at 37 ◻ for 1 h. The formation of mineralized nodules was observed. The results were saved by photography, and the area, quantity and gray scale of mineralized nodules were scanned by Image-Pro Plus 6.0 software for statistical analysis.

### qRT-PCR

The cells were inoculated in 6-well plates and cultured for 24 h for osteogenesis induction. The administration group was given 10 μmol / L TSG, and the control group was added with blank induction medium. Both the administration group and the control group contained 1‰ DMSO. Total RNA was extracted by kit method, and its concentration and purity were determined by ultravioleting-visible spectrophotometer. The RNA concentration was adjusted to 1μg and 1μL was taken as cDNA. Primers and SYBR Green reagent were added to the system. CDNA was amplified by a two-step method on a BIO-RAD CFX96 real-time quantitative PCR instrument. In the pre-denaturation stage, the reaction took place at 95 ◻ for 10 min, with a cycle. The PCR reaction stage was 40 cycles of reaction at 95 ◻ for 15 s, 57 ◻ for 30 s and 72 ◻ for 30 s, respectively. The mRNA expression of the target gene in the drug delivery group and the mRNA expression of the control group were obtained. Internal parameter sequence: GAPDH Forward: TACCCACGGCAAGTTCAACG, Reverse: CACCAGCATCACCCATTTG. Record CT (cyclethreshold) value, through ΔΔ CT = (CT purpose gene-CT inside) treatment group-(CT purpose gene-CT inside) control, computing each group 2 ^−ΔΔCT^.

### Western blot

The cells were inoculated in 6-well plates and cultured for 24 h for osteogenesis induction. The treatment group was given TSG at concentrations of 1, 10 and 50 μmol / L. The blank induction medium was added to the control group, and the culture medium in both the administration group and the control group contained 1‰ DMSO. After 3 days, the total protein was extracted from each well with 300 μL cell lysate (containing PMSF 1 mmol / L). The protein concentration was detected by BCA method. The sample loading buffer containing bromophenol blue was added and denatured at 95 ◻ for 5 min. After 12% SDS-PAGE electrophoresis, the protein was transferred to PVDF membrane. Seal the shaker with 5% skimmed milk powder at room temperature for 1 h. Add KDM5A primary antibody diluted with TBST (Abcam, dilution ratio: 1:1000) and leave at 4 ◻ overnight. The next day, PVDF membrane was washed three times with TBST, and then horseradish peroxidase-labeled secondary antibody diluted with TBST was added. Shake it at room temperature for 2 h. After washing the film for 3 times, the gray value of the protein was detected by an exposure meter.

## Statistical analysis

The results were presented as mean ± standard deviation, SPSS16.0 software was used to process the data, and ANOVA was used to analyze and compare the differences between the multiple groups. T test of students analyzed the comparison between the two groups. P < 0.05 was considered statistically significant.

## Acknowledgements

None.

